# Enculturating a Community of Action (CoA): A qualitative study of health professions educators’ perspectives on teaching with Wikipedia

**DOI:** 10.1101/2021.09.22.461409

**Authors:** Paolo C. Martin, Lauren A. Maggio, Heather Murray, John M. Willinsky

## Abstract

**Purpose:** There is a growing desire for health professions educators to engage learners in more meaningful instruction. Many have tapped Wikipedia to offer an applied approach to engage learners, particularly as it relates to evidence-based medicine (EBM). However, little is known about the benefits and challenges of using Wikipedia as a pedagogical tool from the collective experience of educators who have sought to improve their instructional practice with it. This study aims to uncover and synthesize the perspectives of health professions education (HPE) instructors who have incorporated Wikipedia in their HPE courses.

**Methods:** Applying a constructivist approach, the authors conducted semi-structured interviews with 17 participating HPE instructors who had substantively integrated Wikipedia into their curriculum. Participants were interviewed about their experiences of integrating Wikipedia editing into their courses. Thematic analysis was conducted on resulting transcripts.

**Results:** Authors observed two broad themes among participants’ expressed benefits of teaching with Wikipedia: 1) provides a meaningful instructional alternative that also benefits society and develops learners’ information literacy and EBM skills, and 2) supports learners’ careers and professional identity formation. Identified challenges included: 1) high effort and time, 2) issues with sourcing references, and 3) challenging interactions with skeptics, editors, and students.

**Discussion:** Findings build on known benefits, such as providing a real-world collaborative project that contextualizes students’ learning experiences. They also echo known challenges, such as the resource-intensive nature of teaching with Wikipedia. At the same time, findings extend the current literature by revealing a potential opportunity to approach crowd-sourced information tools, like Wikipedia, as a vehicle to engage and enculturate HPE students within a situated learning context. These findings present implications for HPE programs considering implementing Wikipedia and faculty development needed to help instructors harness crowd-sourced information tools’ pedagogical opportunities as well as anticipate their challenges.

## Introduction

At over 20 years old during the writing of this paper, Wikipedia may be the world’s most viewed medical resource and is an important information tool for health promotion today.^1,2^ Wikipedia’s medical articles have registered over 2 billion views since 2010. In March 2020 alone, when the World Health Organization declared COVID-19 a pandemic, Wikipedia’s English medical articles were visited well over 412 million times.^3^ Wikipedia’s articles are not only visited by the general lay population around the world, including countries with limited access to publicly available health and medical information. Health professions students and clinicians are known to access Wikipedia’s medical articles too.^4–8^ To this point, health professions education (HPE) instructors have used Wikipedia as an educational tool to promote interaction among students and to train them to be critical consumers and contributors to Wikipedia.^9^

A recent inventory of Wikipedia courses aimed at health professions students identified 30 curricular offerings ranging from course assignments to complete courses.^10^ The first reported use of Wikipedia in 2012 was a neuroscience course in which students edited Wikipedia as a course assignment.^11^ Later, Azzam et al.^13^ described the first for-credit medical course fully dedicated to editing and writing medical articles on Wikipedia. Since then, other health professions have reported integrating Wikipedia into their curriculum, including courses in pain management,^13^ pharmacy,^14^ nursing,^15^ and evidence-based medicine.^16^ Despite these offerings, tensions still exist over Wikipedia as a trustworthy information source and whether the academic community should promote its use in their classrooms, with some students and instructors relegated to using Wikipedia as a “guilty secret”.^17^ In the health professions, instructor hesitation over involving a class in contributing to an openly crowd-sourced information resource is understandable. This study speaks to why some instructors believe that this engagement in Wikipedia’s knowledge construction process makes a valuable contribution to HPE.

Because of instructors’ hesitancy about allowing students to use Wikipedia in their classrooms, let alone teach with it, it is important to highlight their insights and perspectives as guidance about how Wikipedia can meet learning objectives within the health professions. Also, as educators seek to apply current technologies with which to engage students, instructors’ perspectives offer unique first-hand accounts about the affordances and challenges of teaching with a public resource designed to engage expert knowledge of medical topics. This could help assist instructors with planning and deciding whether, and to what degree, Wikipedia might be used to impact the experiences of learners in their HPE courses. Because Wikipedia is a collaborative open source platform targeted to the public at large, instructors’ accounts of the pedagogical value of the encyclopedic resource as a teaching tool could inform practical applications of related teaching methods that seek to apply real-world, hands-on, collaborative, activities that might develop civic responsibility, such as service learning approaches or dialogically organized instruction.^18–21^ Towards those aims, this study synthesizes the perspectives of 17 instructors on their motivations to use Wikipedia as a pedagogical tool in HPE, including what instructors perceive to be the value of teaching in this way as well as the practicalities and challenges they have faced in making this curricular choice.

## Methods

Applying a constructivist approach, we conducted semi-structured interviews with 17 participating HPE instructors (July-December, 2020) who integrated Wikipedia into their curriculum. This study was deemed exempt by Stanford University’s institutional review board, (eProtocol #56956).

### Context and Participants

This study recruited 17 instructors in a purposive sampling that drew on an inventory of existing HPE courses that substantively integrated Wikipedia into their teaching approach.^22^ PCM initiated the first round of recruitment to potential participants and scheduled interviews with consenting instructors via email. After the first wave of interviews, the research team convened to review the range of recruited participants and the corresponding data collected to maximize the range of institutional roles and disciplines of HPE instructors recruited in the second wave of emails. If it was revealed during interviews or from course syllabi that an instructor co-taught a course, we also invited those co-instructors to participate.

### Data Collection

PCM conducted and recorded one-on-one interviews with participants via Zoom^23^ that lasted on average one hour, based on a semi-structured interview guide developed by JMW, LAM, and PCM. Interviews were audio recorded with each participant’s permission.

During the interview participants were broadly asked to describe their experiences designing and implementing HPE courses with Wikipedia and were probed for their goals for teaching with Wikipedia, facilitators and barriers they experienced, as well as overall insights and advice they might offer about teaching with Wikipedia (See Appendix A for the interview guide). Upon completion of each interview round, recordings were transcribed and read by PCM and LM to determine if modifications were needed to the interview guide and whether additional recruitment of participants was needed to reach thematic saturation.^24^

### Data Analysis

We used thematic analysis to explore the interview transcripts to derive themes from subsets of interview data related to instructor’s goals and perspectives. Transcripts were coded for themes related to our research questions in batches of 4-6 transcripts over several months. During first cycle coding we applied mostly descriptive and in vivo coding methods and later collapsed initially observed codes into broader categories. A code book was written and revised after each round of coding and was updated as necessary until we could sufficiently identify and describe themes related to instructors’ goals and perspectives about teaching with Wikipedia.

After PCM completed each coding round, PCM, LAM, JMW, and HM met to discuss and resolve coding differences. After resolving those differences, descriptions and examples of observed themes were presented to HM for a check of their resonance as an educator with over three years experience teaching with Wikipedia. We adhered to the Standards for Reporting Qualitative Research checklist for reporting.^25^

### Reflexivity

Our research team included four educators with extensive qualitative methods experience and varying degrees of expertise using Wikipedia as a teaching tool. PCM is an assistant professor of HPE and was a graduate student researcher when this study was conducted. JMW is a professor of education with a background in scholarly communication including its intersection with Wikipedia. LAM is a professor of medicine and HPE who has taught EBM and Wikipedia to medical students.^26^ HM is a professor of emergency medicine and physician who has taught and studied medical education courses with Wikipedia.^27^

## Results

We interviewed 17 instructors representing 13 institutions and 13 HPE courses in the United States (*n*=11), Israel (*n*=1) and Canada (*n*=1). Participants were mostly faculty members (*n*= 8) and librarians (*n*=6). The rest included a lecturer, medical student, and career development specialist. Nine courses fully integrated Wikipedia as a focal aspect of their teaching, while three partially integrated it in the final course project (Table 1). Through our analysis, we observed two themes related to instructors’ perceived benefits and three themes related to perceived challenges of teaching with Wikipedia. These benefits and challenges are presented below. To support these themes, we quote a variety of participants identified by instructor ID (e.g., A).

**Table 1.** Characteristics of 13 health professions education courses that integrated Wikipedia taught by the 17 instructors.

| Course | HPE Program | Instructor IDs | Course Enrollment Size | Elective or Requirement | Course Integration of Wikipedia |
| --- | --- | --- | --- | --- | --- |
| 1 | UME | A,B | 5-37 | ELECT | Full |
| 2 | UME | C | 11 | ELECT | Full |
| 3 | UME | D | 70 | ELECT | Full |
| 4 | UME | E | 77 | REQ | Full |
| 5 | UME | F | 10 | ELECT | Full |
| 6 | UME | G, H, I | 1-4 | ELECT | Full |
| 7 | UME | K | 15 | ELECT | Full |
| 8 | Audiology | L | 10 | REQ | Partial |
| 9 | UME | M, N | 112-120 | REQ | Partial |
| 10 | UME | O | 2-4 | ELECT | Full |
| 11 | GME | P | 8-10 | ELECT | Full |
| 12 | Pharmacology | Q | 10 | REQ | Partial |
| 13 | UME | R | 10 | ELECT | Partial |

Abbreviations: UME = Undergraduate Medical Education, GME = Graduate Medical Education, REQ = Required Course, ELECT = Elective Course, Full = Full course on Wikipedia editing, Partial = Wikipedia editing project embedded into larger course

### Benefits of Teaching with Wikipedia

Although instructors differed in how they came to teach with Wikipedia, they shared two common benefits: 1) to provide a meaningful instructional alternative that would benefit society and develop learners’ information literacy and EBM skills, and 2) to support learners’ careers and professional identity formation.

#### Meaningful Instructional Alternative

Broadly, participants saw Wikipedia as an opportunity to improve learning experiences by situating learning beyond activities that might otherwise be taught in decontextualized ways that threatened student engagement. One instructor, for example, looked to editing Wikipedia as a better way for learners to show “nuance and judgement” compared to multiple choice exams she employed previously (M). Other instructors saw writing Wikipedia medical articles as an alternative assignment that offered more impact than traditional papers, which might only be seen by course instructors. Thus, many instructors hoped that integrating Wikipedia into their teaching would give students a sense of accomplishment, “something they could find value in” (P) and “a feeling like they [students] are part of a larger thing other than themselves” (I).

Wikipedia also imparted, according to two instructors (M & Q), a sense of “gravitas” to their own instructional practices. They believed that Wikipedia did this by developing students’ awareness of social disparities, such as the representation of Wikipedia topics about women in healthcare and women’s health (D). Wikipedia also gave students a chance “to mitigate some of the disparities” by making medical articles on Wikipedia more accurate and up-to-date. This helped society by improving patient access to health information and taking greater ownership over their own healthcare (O).

Some participants also saw integrating Wikipedia as a way to enrich students’ information literacy and EBM skills. By writing Wikipedia articles, students practiced “knowing when to go to what resource” (A) and how to critically appraise the information they found to “enrich a Wikipedia article” (B). Some instructors also hoped students would learn to appreciate the complexity of how Wikipedia worked:

> I wanted them to get down and dirty in the sausage factory, if you will, of Wikipedia, so that they understand the pros and cons of this resource… It’s really helpful for them as practitioners to understand where the evidence is coming from… to click a little bit beyond the text… and be critical about what they’re seeing on the Wikipedia page. (M)

#### Support Students’ Careers and Professional Identity Formation

HPE instructors also hoped that the Wikipedia experience would impact their students’ careers and professional identity formation. For example, some saw Wikipedia as a scholarly activity that could be highlighted on students’ CVs and during residency interviews. One faculty saw their learners’ edits to Wikipedia’s pharmacology articles, which would be seen by others within their discipline, as an opportunity for their learners to “shine” (Q). Many participants also described potential impact on their learners’ professional identities. One faculty hoped Wikipedia would prepare future clinicians who felt “prepared to deliver…services to their patients and industrial clients” (L) and another hoped it would help students, “to have a conversation…on an eye-to-eye level with patients…to receive feedback and give feedback…and to work with people that you might not like” (D). Furthermore, the instructors hoped that by providing learners an in-class opportunity to help society, that their learners would form a sense of responsibility to “do good” with the medical knowledge they encountered. One participant went further to declare:

> My goal, like every educator, wants to transform a student’s life with what they do…this broadens the definition of what it means to be a physician in today’s digitally interconnected era…‘cause if we cede all of Wikipedia authoring to non-health professionals, we are not doing our jobs. (A)

In addition to benefiting students, many participants also found teaching with Wikipedia fulfilling for *themselves*. Those instructors did not directly communicate self-fulfillment as a main benefit of teaching with Wikipedia, but many expressed joy in witnessing students apply knowledge in a new and engaged way. A few instructors also communicated that although teaching with Wikipedia was extra work for them, for which some were not compensated in release time nor in pay, they still decided to teach with it because of the satisfaction it brought them.

### Challenges of Implementing Wikipedia

Although instructors held generally positive views about Wikipedia, they noted three common challenges associated with implementing Wikipedia in their courses: 1) high effort and time, 2) issues with sourcing references, and 3) challenging interactions with skeptics, editors, and students.

#### Effort and Time

Almost all participants only taught HPE courses as a fraction of their total institutional responsibilities. So integrating Wikipedia into their teaching was often a logistical challenge, requiring much effort and time. This challenge led one instructor to question whether she could continue the coordination of many faculty members from specialized disciplines to help advise her 120 students’ Wikipedia writing projects:

> I still wonder like, ‘Will the Wikipedia project live on long term? Is it worthwhile? Or is it just too much work for what the students get out of it?’… It was a boatload of work. I liked doing it…But at the end of the day, I’m not sure. (M)

Other participants described relying on teaching support from librarians, additional faculty members, and WikiEdu staff (volunteer course consultants for instructors) to provide content-area and technical expertise to implement their courses. Another time-consuming challenge for many instructors was learning the technical aspects of teaching with Wikipedia, such as how to edit Wikipedia pages and utilize WikiEdu’s learning management system, and curating Wikipedia training modules necessary for students to begin editing. After course set-up, other reported challenges included delivering synchronous lectures, providing individualized feedback to learners, resolving technical issues, and on some occasions intervening on the students’ behalf to resolve disputes that arose with other Wikipedia editors. While these logistical challenges are shared broadly by most people who teach, teaching with Wikipedia was a “high-touch” endeavor that required an exceptionally high investment of time and resources for many of the instructors in this study.

#### Wikipedia Sourcing Guidelines

According to Wikipedia’s guidelines, medical articles should be based on secondary, but still peer-reviewed, sources of information such as review articles, as well as textbooks. Thus, learners were broadly prohibited from using primary research articles to support their editing of Wikipedia. Most of the instructors understood the general reasoning behind those guidelines. However, many of them also noted the burden those restrictions placed on their courses. In general, the burden resulted from a substantial shift in how learners were used to writing research articles that typically referenced primary sources of information and discouraged from citing secondary sources of information such as textbooks. As one instructor commented, “I think Wikipedia is in the right on this one, but it does take some wrapping your head around mentally” (F). This restriction against primary sources, however, also challenged and frustrated students who wrote articles on less-researched topics, such as rare diseases where there were only primary sources available. Other difficulties noted by participants were finding open access images to upload to support articles, identifying solutions to technical issues that were not covered in WikiEdu training modules, and finding the appropriate articles for students to write about that interested students and filled a need on Wikipedia.

#### Convincing Stakeholders

For many of the instructors, teaching with Wikipedia added a layer of interpersonal challenges. “I don’t like how, when you tell people about it initially, a lot of people will scoff,” one participant (E) described. Skeptics sometimes included students, but were often other HPE colleagues and administrators, whom the instructors had to convince to even be able to teach with Wikipedia. Some instructors noted that skeptics doubted the reliability or scholarly quality of Wikipedia’s content; others, they said, did not value Wikipedia because commonly used resources, such as UpToDate, were perceived as more reliable. Particularly stinging challenges came from the community of Wikipedians who monitored the students’ written contributions and with whom some students had negative interactions, according to some participants: “I found that very frustrating, because there is very little that…I could do about that to basically plead with them not to beat up the students” (B). This issue was especially challenging because the interactions were associated with the removal of students’ contributions from Wikipedia. Sometimes, the disappointment from having their editing deleted was so deep that students did not bother to defend their work to the Wikipedia community, even when they felt they were right. Students also posed interpersonal challenges to some instructors. On rare occasions, some instructors bemoaned those students who did not take their course seriously and fell behind on their assignments. One faculty recounted one exchange, “I went through the trouble of making this course for you. I’m not getting paid to run the course. I do it because I believe…in the power of this course, and I’m helping you transform and grow your identities. If you’re just going to give me shit…those are the most frustrating things” (A).

## Discussion

In this study, we found that instructors’ perspectives echo known benefits of teaching with Wikipedia, especially with regard to teaching EBM and collaborative learning opportunities.^28–30^ Instructors also reported common challenges, like defending Wikipedia to skeptics, motivating students, and its resource-intensive nature.^17,31,32^ For many of the participants, as educators in general, teaching is an art and a passion, but it is often difficult to do well. Indeed as reported by others, suboptimal teaching practices exist in the health professions^33^ and specifically in relation to EBM.^34^ Our findings highlighted a sample of health professions educators’ tenacity to take on those challenges by teaching with Wikipedia in their quest to deliver more meaningful instruction. Participants’ responses extend the current literature by revealing a potential opportunity to approach crowd-sourced information tools like Wikipedia to engage HPE students as a community of action (CoA) within a situated learning context.^35–38^ Within this Wikipedia CoA clinicians are enculturated to be stewards of knowledge with a sense of civic professionalism.^39,40^

### Beyond Motivation and Learning Outcomes

On the broadest level, participants wanted to increase student engagement because they identified issues with prior iterations of their HPE courses, which many described as employing traditional pedagogies such as direct instruction, multiple choice assessments, and summative essays. They were drawn to Wikipedia because it presented an innovative way to structure all or parts of their coursework around a real-world collaborative project with which to contextualize students’ learning experiences. Ostensibly, the hope was to improve learning outcomes, such as clinical content knowledge, and information literacy skills. They also hoped to improve aspects of students’ learning experiences such as their intrinsic motivation^41,42^ within their respective fields.

While concern for student outcomes and learning experiences were important to HPE instructors, our findings also pointed to deeper, less examined motivations for teaching with Wikipedia. Those included their hopes to build a more equitable and just society by improving access to medical information and highlighting the issues of historically excluded and marginalized groups in medical texts. It also included an altruistic desire to help students succeed in their careers and to be critically responsive to social inequities and injustices they encounter in their work.

### Situating Learning in Community of Action

We argue that participants’ reported motivations for teaching with Wikipedia reflect key aspects of Situated Learning Theory (SLT),^37^ particularly in what Wenger and Snyder^36^ might describe as a *community of action* (CoA). According to SLT, novices within a CoA, acquire the skills, values, and beliefs of their respective communities through progressive interactions with knowledgeable others (“elders”) to effect a desired change within or beyond their communities. Teaching with Wikipedia embodied this learning paradigm by capitalizing on an essential activity: the collective, dynamic, and social construction of information through its encyclopedic articles. Indeed, the quality of Wikipedia articles are predicated on the concerted effort of volunteers, who curate, write, edit, and monitor medical articles on Wikipedia. While the interactions of students with the Wikipedia community sometimes produced tense exchanges, by situating students within a CoA, they also apprenticed with other experts, such as their instructors, librarians, seasoned clinicians, and Wikipedia editors, to adopt the skills, values, and beliefs of knowledge construction within their HPE discipline to become actors of social change. In other words, students’ participation in editing Wikipedia was not merely a tool for their own learning, but a means to acquire the experience needed to interact within and outside of their immediate community, and shape their identities as health professionals.

Through a lens of SLT, teaching with Wikipedia might have helped engage students by connecting them to a community larger than themselves, which was dedicated to addressing real-world issues. It also provided role models of what it might mean to be health professionals *and* critical actors of change. Taken together, the participants’ insights about teaching with Wikipedia seem to tell a story about their passion for teaching and how to improve student experiences in the hopes of impacting students’ professional identity formation as civic-minded clinicians. Instructors’ aim to use Wikipedia to advance students’ professional trajectories and enculturate civic professionalism^43^ add to the potential benefits of teaching with Wikipedia. For many health professionals, it honors their hippocratic oath,^44,45^ and promotes the ethical concept of medicine and trustworthiness of the profession.^46^

These findings present implications for HPE programs considering Wikipedia as a teaching tool and faculty development needed to help instructors explore crowd-sourced information tools’ pedagogical opportunities as well as anticipate their challenges. First, they encourage HPE instructors to go beyond a tool’s surface features and consider its potential to offer situated learning experiences, similar to service learning courses, when deciding on adopting tools like Wikipedia to improve instruction. Second, the challenges observed in this study implicate faculty development initiatives needed to destigmatize the use of Wikipedia in HPE courses and training to minimize instructors’ learning curve associated with integrating it into their courses. Lastly, we encourage cross-disciplinary collaboration of experts in information science, online technologies, clinical studies, and Wikipedia editing (e.g., WikiEdu volunteer staff) to guide and support learners’ use of Wikipedia and mediate interactions betweens skeptics and the Wikipedia community.

### Limitations

The findings of this study are not meant to generalize all HPE instructors’ experiences with Wikipedia or draw conclusions about the effects or appropriateness/inappropriateness of using Wikipedia as a tool in any HPE course. We interviewed only a sample of HPE instructors who have taught with Wikipedia, most of them from U.S. undergraduate medical school programs. We also acknowledge that participants’ consent to be part of this study could have largely been related to their positive experiences about teaching with Wikipedia, which they could have been eager to share. As a result we might have missed the voices of HPE instructors who had mostly negative experiences with Wikipedia or unsuccessful attempts at teaching with it. These limitations thus temper the insights we synthesized and recommendations we have expressed about teaching with Wikipedia. Future research on this topic might consider including perspectives from a more diverse sample of HPE instructors and HPE students. Those perspectives might also capture and synthesize their direct recommendations about teaching with and learning from Wikipedia.

## Conclusion

The quality of Wikipedia’s medical articles has been high enough to provide educational benefit to medical students when compared to other commonly approved electronic materials, such as digital textbooks and “UpToDate”.^47^ When considering the rigorous editing process and the engagement of the Wikipedia editing community as well as the high-quality articles produced for the public at large, there are grounds for (re)thinking how health professions education (HPE) students could benefit from contributing to Wikipedia. The idea of teaching with Wikipedia has produced negative reactions,^48^ but we hope that participants’ synthesized perspectives can address hesitancies, and encourage others to explore new and meaningful pedagogical opportunities. The instructors in this study make a compelling case for both Wikipedia’s potential for service learning experiences and rallying students to participate in a critical community of action against social disparities related to knowledge production and information access.

## References

1. Heilman JM, Kemmann E, Bonert M, et al. Wikipedia: a key tool for global public health promotion. J Med Internet Res. 2011;13(1):e14. doi:10.2196/jmir.1589

2. Heilman JM, West AG. Wikipedia and medicine: quantifying readership, editors, and the significance of natural language. J Med Internet Res. 2015;17(3):e62. doi:10.2196/jmir.4069

3. Wikipedia. WikiProject Medicine/Popular pages. https://en.wikipedia.org/w/index.php?title=Wikipedia:WikiProject_Medicine/Popular_pages&oldid=1042725949 Updated September 6, 2021. Accessed September 16, 2021.

4. Beck, J. Doctors’ #1 Source for Healthcare Information: Wikipedia. The Atlantic. March 5, 2014. https://www.theatlantic.com/health/archive/2014/03/doctors-1-source-for-healthcare-information-wikipedia/284206/

5. Aibar E, Lladós-Masllorens J, Meseguer-Artola A, Minguillón J, Lerga M. Wikipedia at university: What faculty think and do about it. The Electronic Library. 2015;33(4):668–683. doi:10.1108/EL-12-2013-0217

6. Knight C, Pryke S. Wikipedia and the University, a case study. Teaching in Higher Education. 2012;17(6):649–659. doi:10.1080/13562517.2012.666734

7. Brox H. The Elephant in the Room: A Place for Wikipedia in Higher Education? Nordlit. 2012;30:143–155. doi:10.7557/13.2377

8. Hughes B, Joshi I, Lemonde H, Wareham J. Junior physician’s use of Web 2.0 for information seeking and medical education: a qualitative study. Int J Med Inform. 2009;78(10):645–655. doi:10.1016/j.ijmedinf.2009.04.008

9. Smith DA. Situating Wikipedia as a health information resource in various contexts: A scoping review. PLoS One. 2020;15(2):e0228786. Published 2020 Feb 18. doi:10.1371/journal.pone.0228786

10. Maggio LA, Willinsky JM, Costello JA, Skinner NA, Martin PC, Dawson JE. Integrating Wikipedia editing into health professions education: a curricular inventory and review of the literature. Perspect Med Educ. 2020;9(6):333–342. doi:10.1007/s40037-020-00620-1

11. Burdo JR. Wikipedia neuroscience stub editing in an introductory undergraduate neuroscience course. J Undergrad Neurosci Educ JUNE Publ FUN Fac Undergrad Neurosci. 2012;11(1):A1–5

12. Azzam A, Bresler D, Leon A, et al. Why Medical Schools Should Embrace Wikipedia: Final-Year Medical Student Contributions to Wikipedia Articles for Academic Credit at One School. Acad Med. 2017;92(2):194–200. doi:10.1097/ACM.0000000000001381

13. Kantarovich D, Vollbrecht HB, Cruz SA, et al. Wikipedia: A Medical Student Educational Project to Edit Wikipedia in Preparation for Practicing Evidence-Based Pain Medicine. J Med Educ Curric Dev. 2020;7:2382120520959691. doi:10.1177/2382120520959691

14. Apollonio DE, Broyde K, Azzam A, De Guia M, Heilman J, Brock T. Pharmacy students can improve access to quality medicines information by editing Wikipedia articles. BMC Med Educ. 2018;18(1):265. doi:10.1186/s12909-018-1375-z

15. Haigh CA. Wikipedia as an evidence source for nursing and healthcare students. Nurse Educ Today. 2011;31(2):135–139. doi:10.1016/j.nedt.2010.05.004

16. Murray H, Walker M, Dawson J, Simper N, Maggio LA. Teaching Evidence-Based Medicine to Medical Students Using Wikipedia as a Platform. Acad Med. 2020;95(3):382–386. doi:10.1097/ACM.0000000000003085

17. Godlee F. Unethical, a guilty secret, and still crazy after all these years. BMJ. 2014;348:g2396. doi:10.1136/bmj.g2396

18. Hunt JB, Bonham C, Jones L. Understanding the goals of service learning and community-based medical education: a systematic review. Acad Med. 2011;86(2):246–251. doi:10.1097/ACM.0b013e3182046481

19. Stewart T, Wubbena ZC. A systematic review of service-learning in medical education: 1998-2012. Teach Learn Med. 2015;27(2):115–122. doi:10.1080/10401334.2015.1011647

20. Kuper A, Boyd VA, Veinot P, et al. A Dialogic Approach to Teaching Person-Centered Care in Graduate Medical Education. J Grad Med Educ. 2019;11(4):460–467. doi:10.4300/JGME-D-19-00085.1

21. Wegerif, R. Dialogic: Education for the Internet Age. England, UK: Routledge; 2013.

22. Maggio LA, Willinsky JM, Costello JA, Skinner NA, Martin PC, Dawson JE. Integrating Wikipedia editing into health professions education: a curricular inventory and review of the literature. Perspect Med Educ. 2020;9(6):333–342. doi:10.1007/s40037-020-00620-1

23. Zoom Video Communications Inc. Security guide. https://d24cgw3uvb9a9h.cloudfront.net/static/81625/doc/Zoom-Security-White-Paper.pdf Published July 2016. Accessed September 16, 2021.

24. Patton MQ. Qualitative Research and Evaluation Methods. 4th Ed. Thousand Oaks, CA: Sage Publications; 2014.

25. O’Brien BC, Harris IB, Beckman TJ, Reed DA, Cook DA. Standards for reporting qualitative research: a synthesis of recommendations. Acad Med. 2014;89(9):1245–1251. doi:10.1097/ACM.0000000000000388

26. Azzam A, Bresler D, Leon A, et al. Why Medical Schools Should Embrace Wikipedia: Final-Year Medical Student Contributions to Wikipedia Articles for Academic Credit at One School. Acad Med. 2017;92(2):194–200. doi:10.1097/ACM.0000000000001381

27. Murray H, Walker M, Dawson J, Simper N, Maggio LA. Teaching Evidence-Based Medicine to Medical Students Using Wikipedia as a Platform. Acad Med. 2020 Mar;95(3):382–386. doi:10.1097/ACM.0000000000003085.

28. Murray H, Walker M, Dawson J, Simper N, Maggio LA. Teaching Evidence-Based Medicine to Medical Students Using Wikipedia as a Platform. Acad Med. 2020;95(3):382–386. doi:10.1097/ACM.0000000000003085

29. Walker M, Murray H, Dawson J, Maggio L. 50: Wikipedia culture and usage: a survey of first year medical students to determine barriers and facilitators. BMJ Evidence-Based Medicine. 2018;23:A25.

30. Smith DA. Situating Wikipedia as a health information resource in various contexts: A scoping review. PLoS One. 2020;15(2):e0228786. Published 2020 Feb 18. doi:10.1371/journal.pone.0228786

31. Bayliss G. Exploring the Cautionary Attitude Toward Wikipedia in Higher Education: Implications for Higher Education Institutions. New Review of Academic Librarianship. 2013;19(1):36–57. doi:10.1080/13614533.2012.740439

32. Dooley PL. Wikipedia and the two-faced professoriate. In: Proceedings of the 6th international symposium on Wikis and open collaboration. 2010;24:1–2. doi:10.1145/1832772.1832803

33. Mehta NB, Hull AL, Young JB, Stoller JK. Just imagine: new paradigms for medical education. Acad Med. 2013;88(10):1418–1423. doi:10.1097/ACM.0b013e3182a36a07

34. Maggio LA, ten Cate O, Chen HC, Irby DM, O’Brien BC. Challenges to Learning Evidence-Based Medicine and Educational Approaches to Meet These Challenges: A Qualitative Study of Selected EBM Curricula in U.S. and Canadian Medical Schools. Acad Med. 2016;91(1):101–106. doi:10.1097/ACM.0000000000000814

35. O’Brien BC, Battista A. Situated learning theory in health professions education research: a scoping review. Adv Health Sci Educ Theory Pract. 2020;25(2):483–509. doi:10.1007/s10459-019-09900-w

36. Wenger EC, Snyder WM. Communities of practice: The organizational frontier. Harvard business review. 2000;78(1):139–146.

37. Lave J, Wenger E. Situated learning: Legitimate peripheral participation. Cambridge, UK: Cambridge university press; 1991.

38. Wenger E. Communities of practice: Learning as a social system. Systems thinker. 1998;9(5):1–5.

39. Boyte HC, Fretz E. Civic professionalism. Journal of Higher Education Outreach and Engagement. 2010;14(2):67–90.

40. Whitcomb ME. Fostering civic professionalism in tomorrow’s doctors. Acad Med. 2005;80(5):413–414. doi:10.1097/00001888-200505000-00001

41. Deci EL, Ryan RM. A motivational approach to self: Integration in personality. In: Dienstbier RA. Nebraska Symposium on Motivation, 1990: Perspectives on motivation. Lincoln, NE: University of Nebraska Press; 1991;237–288.

42. Williams GC, Saizow RB, Ryan RM. The importance of self-determination theory for medical education. Acad Med. 1999;74(9):992–995. doi:10.1097/00001888-199909000-00010

43. Boyte HC, Fretz E. Civic professionalism. Journal of Higher Education Outreach and Engagement. 2010;14(2):67–90.

44. Louis C. Lasagna Oath. Lasagna papers, D.302, Rare Books, Special Collections, and Preservation. River Campus Libraries. Rochester, NY:University of Rochester;1963.

45. Masukume G. Why and how medical schools, peer-reviewed journals, and research funders should promote Wikipedia editing. Studies in Higher Education. 2020;45(5): 984–989. doi:10.1080/03075079.2020.1749796

46. McCullough LB, Coverdale JH, Chervenak FA. Trustworthiness and Professionalism in Academic Medicine. Acad Med. 2020;95(6):828–832. doi:10.1097/ACM.0000000000003248

47. Scaffidi MA, Khan R, Wang C, et al. Comparison of the Impact of Wikipedia, UpToDate, and a Digital Textbook on Short-Term Knowledge Acquisition Among Medical Students: Randomized Controlled Trial of Three Web-Based Resources. JMIR Med Educ. 2017;3(2):e20. doi:10.2196/mededu.8188

48. Jemielniak D. Wikipedia: Why is the common knowledge resource still neglected by academics?. Gigascience. 2019;8(12):giz139. doi:10.1093/gigascience/giz139

